# Production of glycine-derived ammonia as a low-cost and long-distance antibiotic strategy by *Streptomyces*

**DOI:** 10.1101/450833

**Authors:** Mariana Avalos, Paolina Garbeva, Jos M. Raaijmakers, Gilles P. van Wezel

## Abstract

Soil-inhabiting streptomycetes are Nature’s medicine makers, producing over half of all known antibiotics and many other bioactive natural products. However, these bacteria also produce many volatile compounds, and research into these molecules and their role in soil ecology is rapidly gaining momentum. Here we show that streptomycetes have the ability to kill bacteria over long distances via air-borne antibiosis. Our research shows that streptomycetes do so by producing surprisingly high amounts of the low-cost volatile antimicrobial ammonia, which travels over long distances and antagonises both Gram-positive and Gram-negative bacteria. Glycine is required as precursor to produce ammonia, and inactivation of the glycine cleavage system annihilated air-borne antibiosis. As a resistance strategy, *E. coli* cells acquired mutations resulting in reduced expression of the porin master regulator OmpR and its cognate kinase EnvZ, which was just enough to allow them to survive. We further show that ammonia enhances the activity of the more costly canonical antibiotics, suggesting that streptomycetes adopt a low-cost strategy to sensitize competitors for antibiosis over longer distances.

## INTRODUCTION

Volatile Compounds (VC) are small molecules with high vapor pressure and low molecular weight that easily diffuse through air, water or soil(Schmidt et al 2015, Schulz and Dickschat 2007). VCs have a broad activity-spectrum, acting as infochemicals, growth-promoting or inhibiting agents, modulators of quorum sensing and drug resistance or as a carbon-release valve, influencing their neighbor’s behavior and phenotypes such as stress response, colony morphology, biofilm, virulence and pigment production(Audrain et al 2015, Kai et al 2009, Kim et al 2013, Nijland and Burgess 2010, Que et al 2013). In soil, VCs play important roles in inter- and intra-species interactions(Schulz-Bohm et al 2017).

Actinobacteria are one of the largest bacterial phyla present in soil(Barka et al 2016, Cordovez et al 2015). They are known as Nature’s medicine makers (Hopwood 2007b), with the ability to produce bioactive secondary metabolites that work as among others antibiotics, anticancer, antifungal, anthelmintic and immunosuppressant agents(Barka et al 2016, Bérdy 2012, Hopwood 2007a). Streptomycetes alone produce half of all known antibiotics used in the clinic. Streptomycetes are also prolific producers of volatile compounds, often with unknown functions(Citron et al 2015, Schöller et al 2002). In terms of antimicrobial bioactivity of VCs, information is primarily available on their activity as antifungals(Cordovez et al 2015, Wang et al 2013). A rare example of a volatile organic compounds (VOC) with antibacterial activity is the sesquiterpene albaflavenone produced by *Streptomyces albidoflavus* (Gurtler et al 1994).

The natural role of antibiotics is subject to intensive debate. It has been argued that their main function lies in cell to cell communication(Davies 2006). However, antibiotics may well act as weapons, and bioactivity is influenced by social and competitive interactions between strains(Abrudan et al 2015). Interestingly, it is becoming evident that the small inorganic VCs such as hydrogen sulfide (H_2_S) and nitric oxide (NO) play a major role in modulating antibiotic activity and resistance(Avalos et al 2018b). H_2_S production protects bacteria against antibiotics targeting DNA, RNA, protein and cell wall biosynthesis(Shatalin et al 2011). *Bacillus anthracis* produces volatile NO to protect itself against oxidative stress and helps the bacterium to survive in macrophages, thus playing a key role in escaping the host defense(Gusarov and Nudler 2005, Shatalin et al 2008). However, production of NO also directly protects bacteria against a broad spectrum of antibiotics(Gusarov et al 2009, van Sorge et al 2013). Ammonia induced resistance to tetracycline, by increasing the level of polyamines which leads to a modification of membrane permeability(Bernier et al 2011). There is also some experimental evidence that suggests that VCs may affect membrane integrity(Fadli et al 2014, Yung et al 2016), which in turn may make the cells more susceptible to other cell-damaging compounds, such as antibiotics. The lack of information makes it hard to mimic the biological effect and more so to pinpoint the responsible molecules of such activity.

In this study we show for the first time that *Streptomyces* can produce surprisingly high levels of ammonia that affect surrounding bacteria even over long distances. The concentrations accumulated away from the colonies are so high that the ammonia acts as an antibiotic, inhibiting the growth of *Escherichia coli* and *Bacillus subtilis*. We also show that the production of ammonia in *Streptomyces* can be controlled by varying growth conditions, and depends on the glycine cleavage system. *E. coli* cells gain resistance against the ammonia by reducing the expression of the two-component system OmpR-EnvZ, thereby counteracting passage through the outer membrane porins.

## MATERIALS AND METHODS

### Strains, media, culture conditions and antimicrobial assays

Strains used in this study are listed in Table S1. The *Streptomyces* strains were grown on Soy Flour Mannitol (SFM) agar plates to prepare spore stocks. *Escherichia coli* strain AS19-RlmA-(Liu and Douthwaite 2002) and *B. subtilis* 168(Barbe et al 2009) were used as test microorganisms and grown on Luria-Bertani (LB) agar plates.

Volatile antimicrobial assays were performed using a petri dish with two compartments, one filled with SFM media for *Streptomyces* growth and the second one with LB +/- TES buffer 50-100 mM. *Streptomyces* strains were streaked on the SFM side and allowed to grow for 5 days after which, *E. coli* or *B. subtilis* were inoculated on the LB side using a concentration of 10^4^ and 10^3^ cfu/mL respectively.

### Collection and analysis of VCs

VCs produced by *Streptomyces* monocultures grown on SFM agar were collected using a glass petridish designed for trapping of the volatile headspace(Garbeva et al 2014). The lid of the glass petridish contains an outlet specially designed to hold a stainless steel column packed with 200 mg Tenax^®^ TA 60/80 material (CAMSCO, Houston, TX, USA). Samples were taken in triplicates from day 3 to day 5 of growth; after that, the Tenax steel traps were sealed and stored at 4°C until GC-Q-TOF analysis.

Trapped volatiles were desorbed using an automated thermodesorption unit (model UnityTD-100, Markes International Ltd., United Kingdom) at 210°C for 12 min (Helium flow 50 ml/min) and trapped on a cold trap at -10°C. The trapped volatiles were introduced into the GC-QTOF (model Agilent 7890B GC and the Agilent 7200A QTOF, USA) by heating the cold trap for 3 min to 280°C. A 30 × 0.25 mm ID RXI-5MS column with a film thickness of 0.25 μm was used (Restek 13424-6850, USA). Temperature program used was as follows: 39°C for 2 min, from 39 to 95°C at 3,5 °C/min, then to 165°C at 6°C/min, to 250°C at 15°C/min and finally to 300°C at 40°C/min, hold 20 min. The VCs were detected by the mass spectrometer (MS) operating at 70 eV in EI mode. MS spectra were extracted with MassHunter Qualitative Analysis Software V B.06.00 Build 6.0.633.0 (Agilent Technologies, USA) using the GC-Q-TOF qualitative analysis module. MS spectra were exported as mzData files for further processing in MZmine. The files were imported to MZmine V2.14.2(Pluskal et al 2010) and compounds were identified via their mass spectra using deconvolution function (Local-Maximum algorithm) in combination with two mass-spectral-libraries: NIST 2014 V2.20 (National Institute of Standards and Technology, USA http://www.nist.gov) and Wiley 9th edition mass spectral libraries and by their linear retention indexes (LRI). The LRI values were calculated using an alkane calibration mix before the measurements in combination with AMDIS 2.72 (National Institute of Standards and Technology, USA). The calculated LRI were compared with those found in the NIST and in the in-house NIOO-KNAW LRI database. After deconvolution and mass identification peak lists containing the mass features of each treatment (MZ-value/Retention time and the peak intensity) were created and exported as CSV files for statistical processing via MetaboAnalyst V3.0 (www.metaboanalyst.ca; (Xia et al 2015)).

### pH change, ammonia determination and toxicity

Change in pH of the growth media was determined by the addition of phenol red indicator (0.002%). Pictures were taken after 0, 3 and 5 days of incubation next to *Streptomyces* biomass. For the ammonia test, *Streptomyces* strains were grown for 5 days on SFM agar using the two-compartment petri-dish, whereby the other half of the plate was left empty. After 5 days, the ammonia was determined using the Quantofix^®^ ammonium test kit. Pictures were recorded to obtain a qualitative measurement of ammonia production from each strain.

Quantification of ammonia accumulation inside the LB agar was determined by extracting the liquid from the LB agar by centrifugation. For this, centrifuge tube filters were used (spin-X^®^ 0.22 μm cellulose acetate, Corning Inc. USA), 1 cm^2^ of agar was put inside the filter tube and centrifuge at 13,000 rpm for 20 min. The eluate (∼200 μL) was used to quantify the ammonium concentration in comparison to a standard curve. The standard curve was made with LB agar containing 0-50 mM concentrations of ammonia. Ammonia solution (25% in H_2_O, J.T. Baker 6051) was used as source of ammonia. The liquid was extracted from the agar the same way as described before and used together with the Quantofix^®^ ammonium kit to obtain a semi-quantitative measure of ammonia accumulation inside the agar.

To determine the toxicity of ammonia, *E. coli* and *B. subtilis* were incubated in the automated Bioscreen C (Lab systems Helsinki, Finland) in the presence of increasing concentrations of ammonia. Each dilution was prepared in LB containing an inoculum of 10^5^cfu/mL + different volumes of ammonia solution (J.T. Baker) to give the following final concentrations: 1,5,10,15,16,17,18,20, 25, 30, 40 and 50 mM. The final working volume in each well of the honeycomb was 100 μL. Cultures were incubated at 37°C overnight with continuous shaking. O.D. measurements (wideband) were taken every 30 min for 20 h. The data and growth curves were calculated from triplicates.

### HCN determination

To detect hydrogen cyanide in the headspace of *Streptomyces* growth we used a method adapted from Castric and Castric(Castric and Castric 1983). For this, Whatman™ paper was soaked in suspension containing 5 mg/ml of copper(II) ethyl acetoacetate and 4,4’-methylenebis-(N, N-dimethylaniline) (Sigma-Aldrich, USA) dissolved in chloroform and allowed to dry protected from light. The filter paper was placed next to Streptomyces pre-grown for 2 days. *Pseudomonas donghuensis* P482 was used as positive control. Strains were incubated at 30°C until blue coloration of the filter paper was evident.

### Whole genome sequencing

Genome sequencing of *E. coli* AS19-RlmA^-^ (Avalos et al 2018a) and its mutant ARM9 was performed using Illumina HiSEQ and PacBio RfavaS at Baseclear BV, Leiden (The Netherlands). Paired-end sequence reads were generated using the Illumina HiSeq 2000 system and mapping the individual reads against the reference genome of *E. coli* B str. REL606. The contigs were placed into superscaffolds based on the alignment of the PacBio CLC reads. Alignment was performed with BLASR(Chaisson and Tesler 2012). Genome annotation was performed using the Baseclear annotation pipeline based on the Prokaryotic Genome Annotation System (http://vicbioinformatics.com). Variant detection was performed using the CLC genomics workbench version 6.5. The initial list of variants was filtered using the Phred quality score and false positives were reduced by setting the minimum variant frequency to 70% and the minimum number of reads that should cover a position was set to 10. Relevant mutations were confirmed by PCR analysis. The genome of *E. coli* AS19-RlmA^-^ has been published, with accession number CP027430 (Avalos et al 2018a).

### Genetic complementation of *ompR* and *envZ*

*E. coli* strain AS19-RlmA^-^ suppressor mutant ARM9 was complemented by inserting the *ompR* or *envZ* genes in pCA24N from the ASKA collection(Kitagawa et al 2005). Cells of suppressor mutant ARM9 containing the plasmid were inoculated in LB + Chloramphenicol (25 μg/mL) with or without IPTG 0.1 mM for induction of the gene expression.

### RNA sequencing

For RNA extraction *E. coli* cells were grown to an O.D._600_ of 0.5, RNA Protect Bacteria Reagent (Qiagen Cat No. 76506) was added according to manufacturer instructions. Cells were pelleted and re-suspended in boiling 2% SDS + 16 mM EDTA followed by extraction with Phenol:chloroform:Isoamyl alcohol (25:24:1) pH 6.6. (VWR Prolabo 436734C). Aqueous phase was precipitated with 3M sodium acetate pH 5.2 and pure ethanol, washed with 70% Ethanol and re-suspended in RNAse-free water. DNA was removed using 5 units of DNAseI (Fermentas #EN0521) with further purification using again phenol:chloroform:isoamyl alcohol and precipitation with sodium acetate and ethanol. The final pellet was dissolved in RNase free water.

RNA sequencing and analysis was performed by Baseclear BV (Leiden, The Netherlands). Ribosomal RNA was subsequently removed with a Ribo-Zero kit (Epicenter) and the remaining RNA used as input for the Illumina TruSeq RNA-seq library preparation. Once fragmented and converted into double strand cDNA, the fragments (about 100-200 bp) were ligated with DNA adapters at both ends and amplified via PCR. The resulting library was then sequenced using an Illumina Sequencer. The FASTQ sequence reads were generated using the Illumina Casava pipeline version 1.8.3. Initial quality assessment was based on data passing the Illumina Chastity filtering. Subsequently, reads containing adapters and/or PhiX control signals were removed using an in-house filtering protocol. The second quality assessment was based on the remaining reads using the FASTQC quality control tool version 0.10.0.

For the RNA-Seq analysis the quality of the FASTQ sequences was enhanced by trimming off low-quality bases using the “Trim sequences” option present in CLC Genomics Workbench Version 6.0.4 (QIAGEN, Bioinformatics). The quality-filtered sequence reads were used for further analysis with CLC Genomics Workbench. First an alignment against the reference and calculation of the transcript levels was performed using the “RNA-Seq” option. Subsequent comparison of transcript levels between strains and statistical analysis was done with the “Expression analysis” option, calculating so-called RPKM values. These are defined as the reads per kilobase per million mapped reads(Mortazavi et al 2008) and normalizes for the difference in the number of mapped reads between samples and for transcript length. The RNAseq data has been submitted to the GEO (Gene Expression Omnibus) from NCBI (National Biotechnology Center Information) with GEO accession number GSE111370.

### Synergism of *Streptomyces* AB-VCs with soluble antibiotics

Synergistic assays were performed using a petri dish with two compartments, one filled with SFM media for *Streptomyces* growth and the second one with LB. *Streptomyces* strains were streaked on the SFM side and allowed to grow for 5 days at 30°C after which, *E. coli* or *B. subtilis* were inoculated on the LB side using a culture grown to an O.D. = 0.5. 100 μL were streaked on the LB side. Afterwards, 6 mm filter discs (Whatman™) were placed on top of the LB and 10 μL of each antibiotic spotted on the filter disc. Two different concentrations per antibiotic were tested: ampicillin 500, 31 μg/mL; erythromycin 250, 31 μg/mL; kanamycin 1000, 500 μg/mL; tylosin 500, 62 μg/mL; actinomycin 500, 62 μg/mL; spectinomycin 1000, 500 μg/mL; streptomycin 500, 62 μg/mL. Plates were incubated at 37°C overnight and pictures recorded after 20 h of growth.

## RESULTS AND DISCUSSION

### VCs as bioactive agents in long-distance antibiosis

Since streptomycetes actively produce antifungal VCs, we set out to investigate if also antibacterial VCs could be produced by these bacteria. After all, this would give them a competitive advantage by inhibiting other bacteria at longer distances. To investigate this, streptomycetes were grown physically separated from indicator/target bacteria by a polystyrene barrier. Air-borne VCs can pass over the barrier, but it does not allow passage of canonical (soluble) antibiotics. As indicator strains we used *Bacillus subtilis* and *Escherichia coli* strain AS19-RlmA^-^ (referred to as *E. coli* ASD19 from now on). The latter has known antibiotic sensitivity(Liu and Douthwaite 2002). Interestingly, the streptomycetes showed varying volatile activity against *E. coli* ASD19; the indicator strain failed to grow adjacent to *Streptomyces* sp. MBT11 or *S. venezuelae*, but grew normally next to *S. coelicolor*, *S. lividans* and *S. griseus* (Figure 1A). *B. subtilis* was not inhibited by any of the former strains.

**Figure 1.**
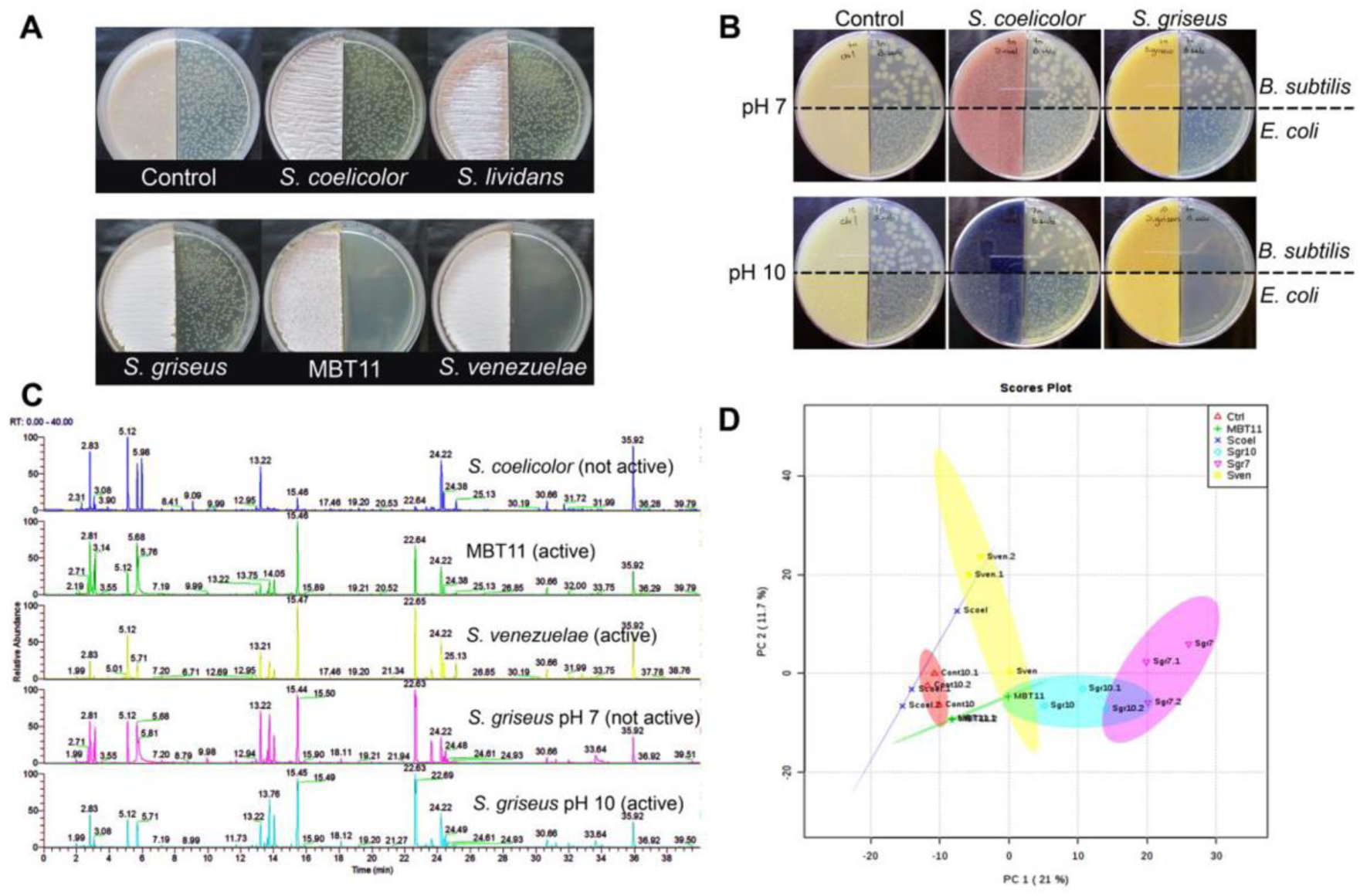
Bioactivity and metabolomic analysis of VCs released by streptomycetes. **A.** Bioactivity of VCs released by selected *Streptomyces* strains against *E. coli* strain ASD19. **B.** Volatile antibiotic activity of different *Streptomyces* strains grown at pH 7 or pH10, the latter by addition of a glycine/NaOH buffer; *E. coli* strain ASD19 was the indicator strain. **C.** Comparison of GC-chromatogram of VCs from bioactive and non-bioactive strains. **D**. PCA-2D Plot showing the VC-grouping of the different strains. No clear separation between VCs produced by different *Streptomyces* strains is seen.

We then wanted to assess whether the production of antimicrobial volatile compounds (AMVCs) could be elicited by varying the growth conditions. We previously showed that growth at pH 10, *N*-acetylglucosamine, starch or yeast extract pleiotropically enhanced the production of antibiotics in many *Streptomyces* species(Zhu et al 2014). Interestingly, in contrast to a neutral pH, when glycine/NaOH buffer was added to raise the pH to 10, *S. griseus* produced VCs that completely inhibited growth of *E. coli* and *B. subtilis* (Figure 1B). AMVC production by *Streptomyces* species MBT11 was also enhanced by growth in the presence of the buffer system. In contrast, *S. coelicolor* failed to produce AMVCs under any of the conditions tested (Table S2).

The induction of AMVCs by *S. griseus* when grown on the glycine buffer offered an ideal system to elucidate the nature of the bioactive molecules by statistical methods. Correlation between bioactivity and metabolic profiles allows efficient reduction of candidate molecules (Gubbens et al 2014, Wu et al 2015). Therefore, GC-Q-TOF-based metabolomics was performed to compare the VC profiles of *S. coelicolor*, *Streptomyces* sp. MBT11, *S. venezuelae* and *S. griseus* (the latter grown with and without glycine buffer pH 10). Despite the antimicrobial activity, no volatile organic compound (VOC) was detected that correlated statistically to the bioactivity, nor did we see any significant difference between the metabolome profiles of *S. griseus* grown with or without the glycine buffer (Figure 1C, D). Some of the mass features suggested differential production of 2-methylisoborneol (2-MIB) and 2-methylenebornane by the active strains compared to the non-active strains. However, mutants of *S. griseus* that lacked either or even all of the terpene cyclases, still retained their volatile antibacterial activity (data not shown).

### The main inhibitory molecule is ammonia

VCs may induce a change in pH away from colonies(Jones et al 2017, Letoffe et al 2014), and we therefore wondered whether the VC bioactivity was accompanied by a pH change on the receiver side. To assess this, we used phenol red, which changes from pale orange to bright pink when the media becomes alkaline. Interestingly, a gradual increase was seen in the pH caused by VCs produced by *S. venezuelae* or by *Streptomyces* sp. MBT11 that produce AMVCs, but not by the non-producing *S. coelicolor*. Initially, a pH increase was seen close to the *Streptomyces* biomass, and after 5 days the receiver side had turned completely pink (Figure 2A). At that point, the pH had increased to around 8.5. The pH increase correlated fully to the antibiosis, with growth inhibition of *E. coli* close to the *Streptomyces* biomass after 3 days, while after 5 days the growth of *E. coli* was fully inhibited (Figure 2B). Further in support of pH-dependent growth inhibition, *E. coli* grew normally when the media was buffered with 50 mM TES (pH 7). However, the pH itself was not the cause of the inhibition, since the *E. coli* cells grew apparently normal on media adjusted to pH 9 (Figure 2C). Also, we previously showed that even at pH 10 antibiotic susceptibility is similar to growth at pH 7(Zhu et al 2014).

**Figure 2.**
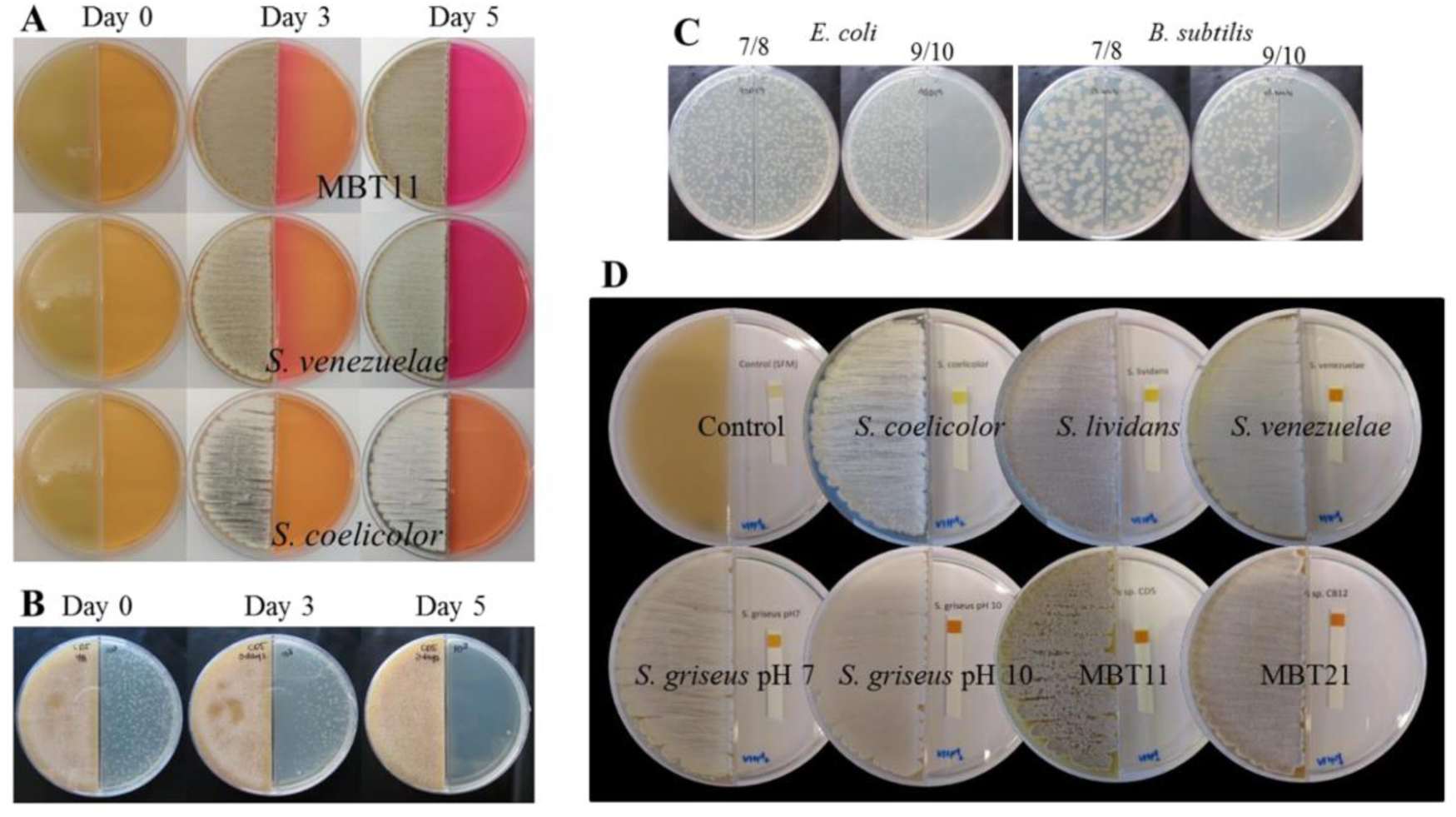
pH increase is caused by high ammonia production. **A.** pH change illustrated by the color change of the indicator (Phenol red 0.002%). LB medium alkalization after 3 and 5 days of growth of *Streptomyces* sp. MBT11 and *S. venezuelae*, no alkalization was caused by VCs produced by *S. coelicolor*. **B.** *Streptomyces* sp. MBT11 antimicrobial VCs production curve against *E. coli* strain ASD19. **C.** *E. coli* strain ASD19 and *B. subtilis* growth under different pH adjusted with glycine/NaOH buffer. **D.** NH3 emission. Test strips on the right compartment show the production of NH3 by *Streptomyces* strains. *S. coelicolor* and *S. lividans* (~10 mg/L); *S. venezuelae* (~100 mg/L); *Streptomyces* sp. MBT11 (~100 mg/L); Control: SFM media (0 mg/L). Concentrations are estimated according to the color chart indicator from the Quantofix^®^ ammonium detection Kit.

Ammonia and trimethylamine are VCs known to induce a pH change(Bernier et al 2011, Čepl JJ 2010, Jones et al 2017, Letoffe et al 2014). We also tested production of hydrogen cyanide (HCN), a known AMVC produced by, among others, rhizospheric streptomycetes(Anwar et al 2016); however, none of the strains produced detectable amounts of HCN (Figure S1). Furthermore, under our growth conditions, TMA was not detected in the headspace of the *Streptomyces* strains (Figure S2). Ammonia production was determined using the Quantofix^®^ Ammonium detection kit. Interestingly, antibiosis by the streptomycetes fully correlated to an increase in ammonia production (Figure 2D). To determine how much ammonia was accumulated, the *Streptomyces* strains were grown for 5 days, and the agar on the receiver side extracted. Agar containing different concentrations of ammonia was used to create a standard curve. *S. coelicolor*, *S. lividans* and *S. griseus* (pH 7) accumulated only 2-5 mM ammonia, while the growth-inhibiting *Streptomyces* sp. MBT11, *S. venezuelae* and *S. griseus* (the latter only with added Gly/NaOH buffer pH 10) had accumulated between 15-30 mM ammonia (Figure 3A).

**Figure 3.**
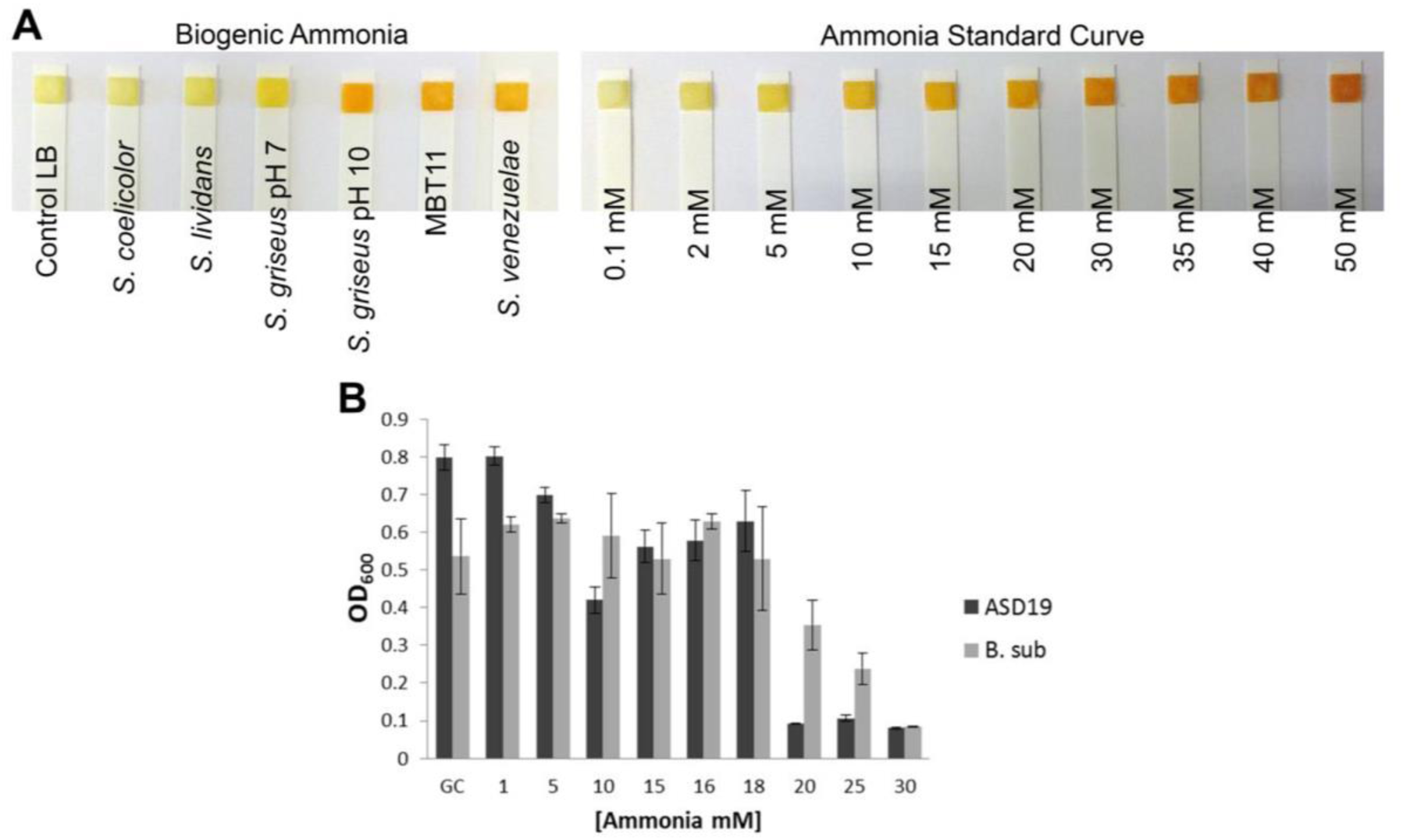
Bioactivity is caused by ammonia. **A.** Ammonia quantification from LB agar extracts exposed to *Streptomyces* VCs (left). Ammonia standard curve from LB agar extract (right). **B.** Growth of *E. coli* ASD19 (black) and *B. subtilis* (gray) under different concentrations of ammonia.

This strongly suggested that ammonia was the AMVC produced by the strains. Indeed, *E. coli* failed to grow on media with 20 mM ammonia or higher, while growth of *B. subtilis* was inhibited by ammonia concentrations above 30 mM (Figure 3B). These values are within the range produced by the strains causing volatile antibiosis.

### Ammonia is derived from glycine cleavage

We then wondered if ammonia was generated from glycine metabolism, because a glycine/NaOH buffer was used to set the pH. A major pathway for the catabolism of glycine is the glycine cleavage system (GCV) that converts glycine into CO_2_, ammonia and a methylene group that is transferred to tetrahydrofolate (THF) to form N_5_,N_10_-methylene-THF(Kikuchi et al 2008, Tezuka and Ohnishi 2014). Importantly, when *S. griseus* was grown on SFM agar containing just glycine at concentrations as low as 0.1% (w/v), this time without increasing the pH, the strain still fully inhibited the growth of *B. subtilis* and *E. coli* (Figure 4A). We then also grew *S. coelicolor* on SFM agar with increasing concentrations of glycine. Interestingly, at concentrations of 1% (w/v) glycine or higher, also *S. coelicolor* fully inhibited the indicator cells. This suggests that at sufficiently high concentrations of glycine, all streptomycetes may produce so much ammonia that it inhibits the growth of other bacteria.

**Figure 4.**
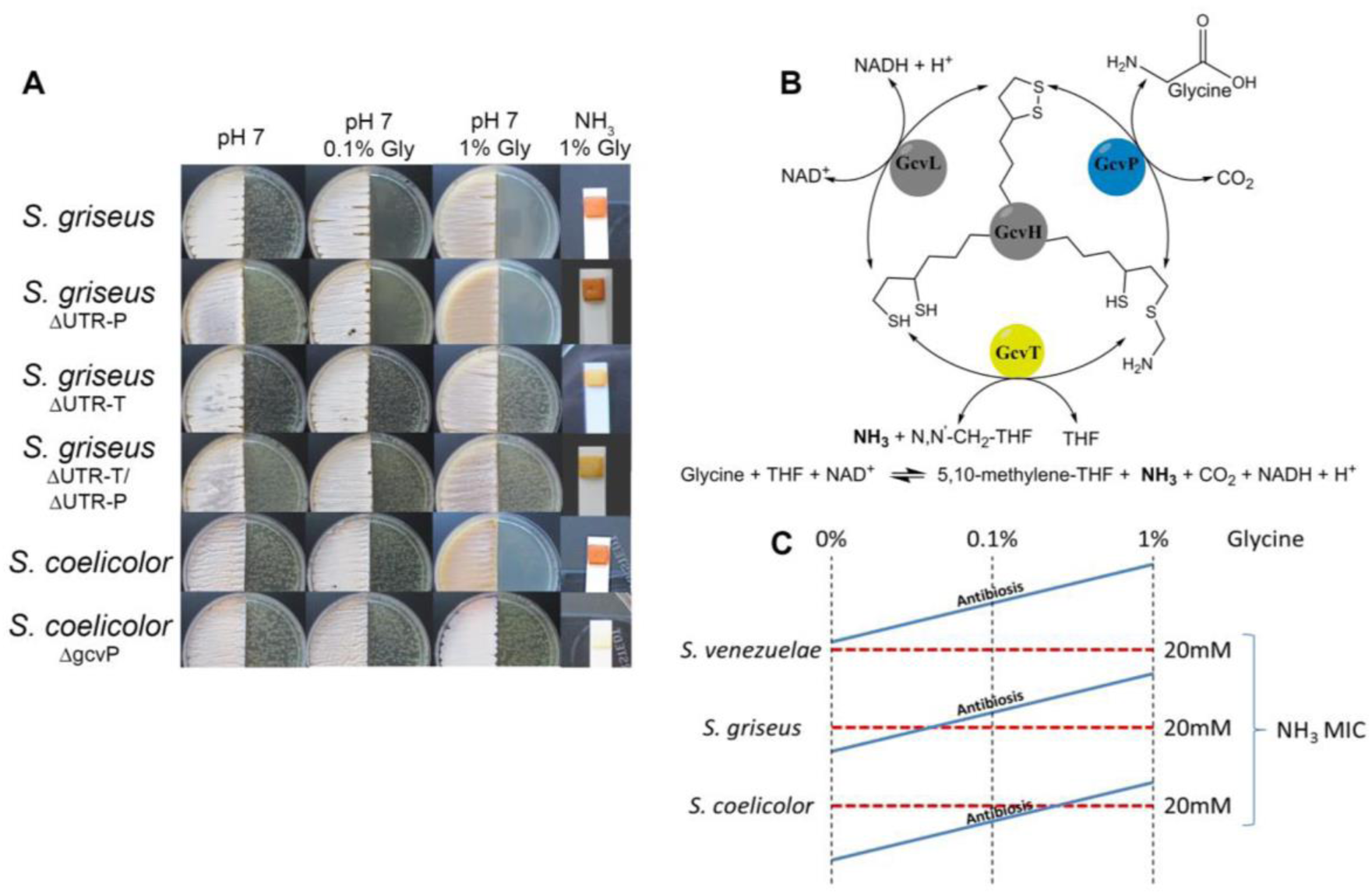
Bioactivity is caused by ammonia in a glycine cleavage-dependent manner. **A.** Volatile activity and ammonia production by *S. griseus*, *S. griseus* glycine cleavage mutant *gcv* P (ΔUTR-P), *S. griseus* glycine cleavage mutant *gcv* T (ΔUTR-T), *S. griseus* glycine cleavage double mutant *gcvT-gcvP* (ΔUTR-T/ΔUTR-P), *S. coelicolor* and the *S. coelicolor* glycine cleavage system mutant Δ*gcv* P. **B.** Scheme representation of the reactions carried by the glycine cleavage (GCV) system, consisting of: pyridoxal phosphate-containing glycine decarboxylase GcvP (blue); THF-dependent aminomethyltransferase GcvT (yellow); dihydrolipoamide dehydrogenase GcvL; and lipoic acid-containing carrier protein GcvH. **C.** Illustration of the induction of volatile antibiosis in different *Streptomyces* strains when increasing concentrations of glycine are added.

We then tested the direct involvement of the GCV system(Tezuka and Ohnishi 2014), which consists of three enzymes (GcvL, GcvP, GcvT) and a carrier protein: GcvH (Figure 4B). Conversely, *gcvP* mutants of *S. coelicolor*(Zhang 2015) and *gcvT* mutants of *S. griseus*(Tezuka and Ohnishi 2014) were unable to produce ihibiting amounts of ammonia, even when grown on high amounts of glycine. A mutant of *S. griseus* lacking the 5’UTR of *gcvP* (Tezuka and Ohnishi 2014) still produced sufficient ammonia to inhibit the growth of *E. coli* cells, but this was annihilated by the additional deletion of *gcvT* (Figure 4A). Taken together, this strongly suggests that in both *S. coelicolor* and *S. griseus*, volatile ammonia is primarily derived from the GCV system, and as expected, GcvT is the key enzyme responsible for the production of ammonia from glycine. Inactivation of *gcvP* is also sufficient to block volatile ammonia production in *S. coelicolor*, fully in line with the idea that the key system is GCV, while in *S. griseus* GcvP can be by-passed by another (yet unknown) enzyme.

A major difference between *S. coelicolor* and *S. griseus* on the one had, and *S. venezuelae* and *Streptomyces* sp. MBT11 on the other, is that the latter two strains do not require any added glycine to produce levels of ammonia above the MIC. Ammonia may be derived from various metabolic enzymes, such as ammonia lyases, deaminases, deiminases and pyridoxamine phosphate oxidases. We are currently performing a large-scale phylogenomics and mutational analysis to identify the gene(s) that are responsible for the overproduction of ammonia in these strains.

### OmpR is key to ammonia resistance

To obtain more insights into the cellular response of *E. coli* cells to ammonia, we selected for spontaneous ammonia-resistant mutants. After two days of growth next to *Streptomyces* sp. MBT11, several colonies appeared that were able to withstand the accumulated ammonia and likely had sustained one or more suppressor mutations. Four of these colonies were analyzed further, which showed different levels of resistance as indicated by the colony size (Figure S3A). Of these, suppressor mutant ARM9 was selected for its high resistance (Figure 5A left). Strain ARM9 was reproducibly more resistant to ammonia than its parent, with MICs of 25 mM and 20 mM, respectively. This is a highly significant difference, as we previously showed that 20 mM is precisely the tipping point for ammonia sensitivity of *E. coli* (Figure 5B).

**Figure 5.**
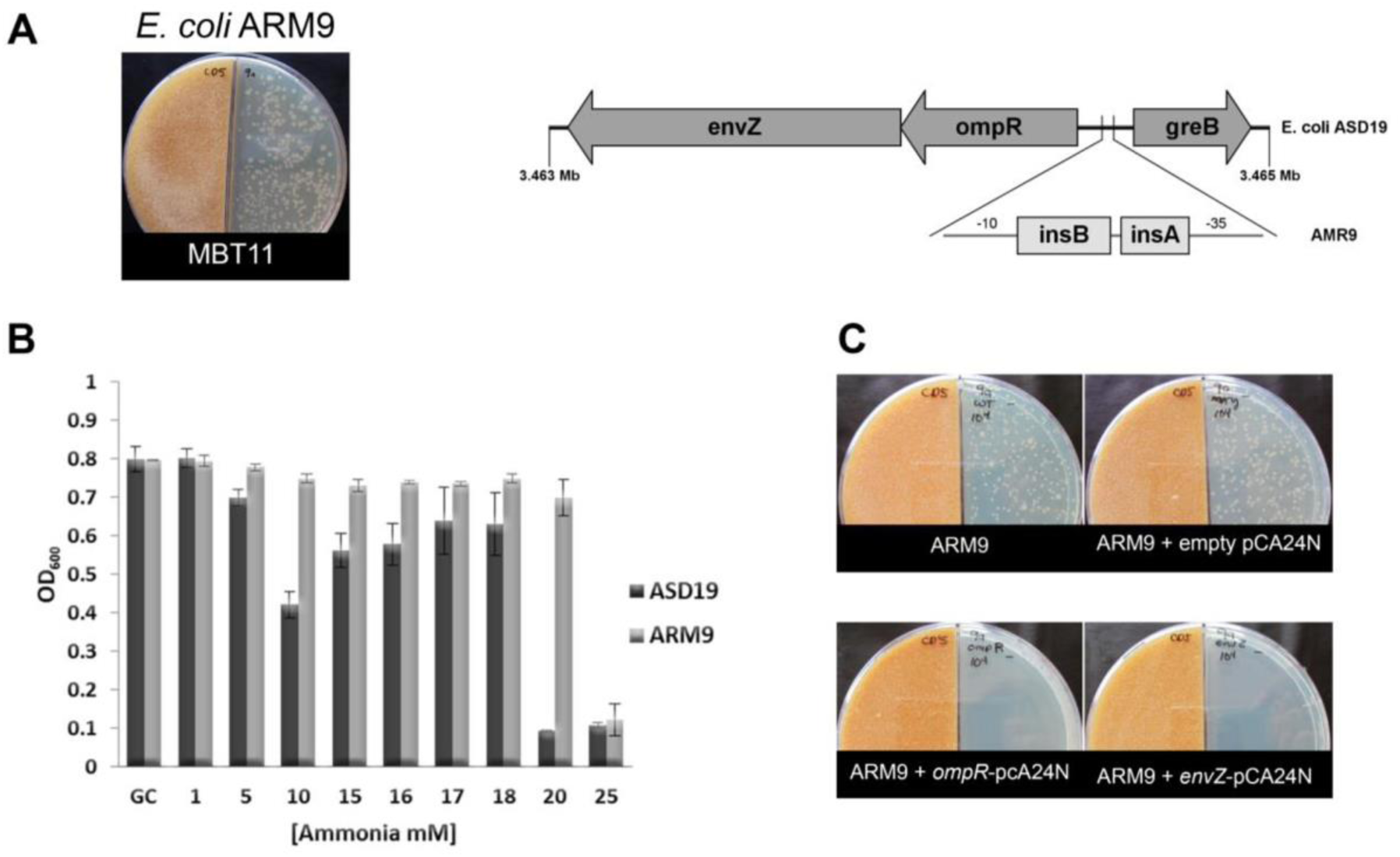
Insertion sequences in the *E. coli ompR-envZ* promoter govern resistance to ammonia. **A.** (left) spontaneous suppressor mutant ARM9 derived from *E. coli* strain ASD19 growing under the presence of volatile compounds produced by *Streptomyces* sp. MBT11; (right) visualization of insertion sequences in-between the -10 and -35 sequences of the *ompR/envZ* promoter. **B.** Growth of *E. coli* strain ASD19 (black) and *E. coli* strain ASD19 suppressor mutant ARM9 (gray) under the presence of different concentrations of ammonia. **C.** Growth of suppressor mutant ARM9 and transformants harbouring either empty plasmid pCA24N, plasmid *ompR*- pCA24N (expressing *ompR*), or plasmid *envZ*-pCA24N (expressing *envZ*). Note that introduction of a plasmid expressing either *envZ* or *ompR* makes ARM9 sensitive again to ammonia.

Under low availability of nitrogen, the AmtB transporter facilitates the intake of ammonium inside the cell(Conroy et al 2007, Wirén and Merrick 2004). Our conditions include high concentrations of ammonia, therefore we hypothesized that a mechanism other than the AmtB channel would be involved in the resistance towards ammonia. To identify the nature of the mutation(s) sustained by ARM9, its genome sequence was compared to that of its parent *E. coli* ASD19 (Table S3). In total 658 mutations were found by single nucleotide permutation (SNP) analysis, of which 198 gave rise to amino acid changes or insertions or deletions. However, one change immediately stood out, namely the introduction of two insertion elements (insA_31 and insB_31) in-between the -35 and -10 consensus sequences of the promoter for *ompR*-envZ, which encode the two-component system (TCS) consisting of response regulator OmpR and sensory kinase EnvZ (Figure 5A right). This TCS is involved in osmoregulation in response to environmental signals(Nikaido 2003) and regulates the expression of outer membrane porins OmpF and OmpC. Importantly, these are known to be involved in antibiotic resistance regulated by osmotic pressure and pH(Fernandez and Hancock 2012), and to reduce the responsiveness of *E. coli* cells to VCs(Kim et al 2013).

### Reduced transcription of the *ompR-envZ* operon is the cause of ammonia resistance

Considering the location right in the middle of the promoter, we expected that the IS elements in the *ompR-envZ* promoter reduced the transcription of these crucial TCS genes. To establish the transcriptional consequences of the IS insertion into the *ompR-envZ* promoter region, RNAseq was performed on *E. coli* ASD19 and its suppressor mutant ARM9 grown in LB media until mid-exponential phase (OD_600_ 0.5), and the global transcription profiles compared (see Table S4 for the full dataset). Table 1 shows genes highly up/down regulated as a result of a clustering analysis using a cut-off value of a fold change +/− 2.0. These data confirm the downregulation of *ompR* and *envZ* genes and other related genetic elements like *omrA*, a small mRNA that negatively regulates *ompR* expression. Additionally, genes involved in amino-acid metabolism were down regulated, including the *astABCE* gene cluster involved in the ammonia-producing arginine catabolic pathway, *aspA* that is involved in the conversion of L- aspartate into fumarate and ammonia, and *tnaC* for catabolism of tryptophan, which again releases ammonia.

**Table 1.**
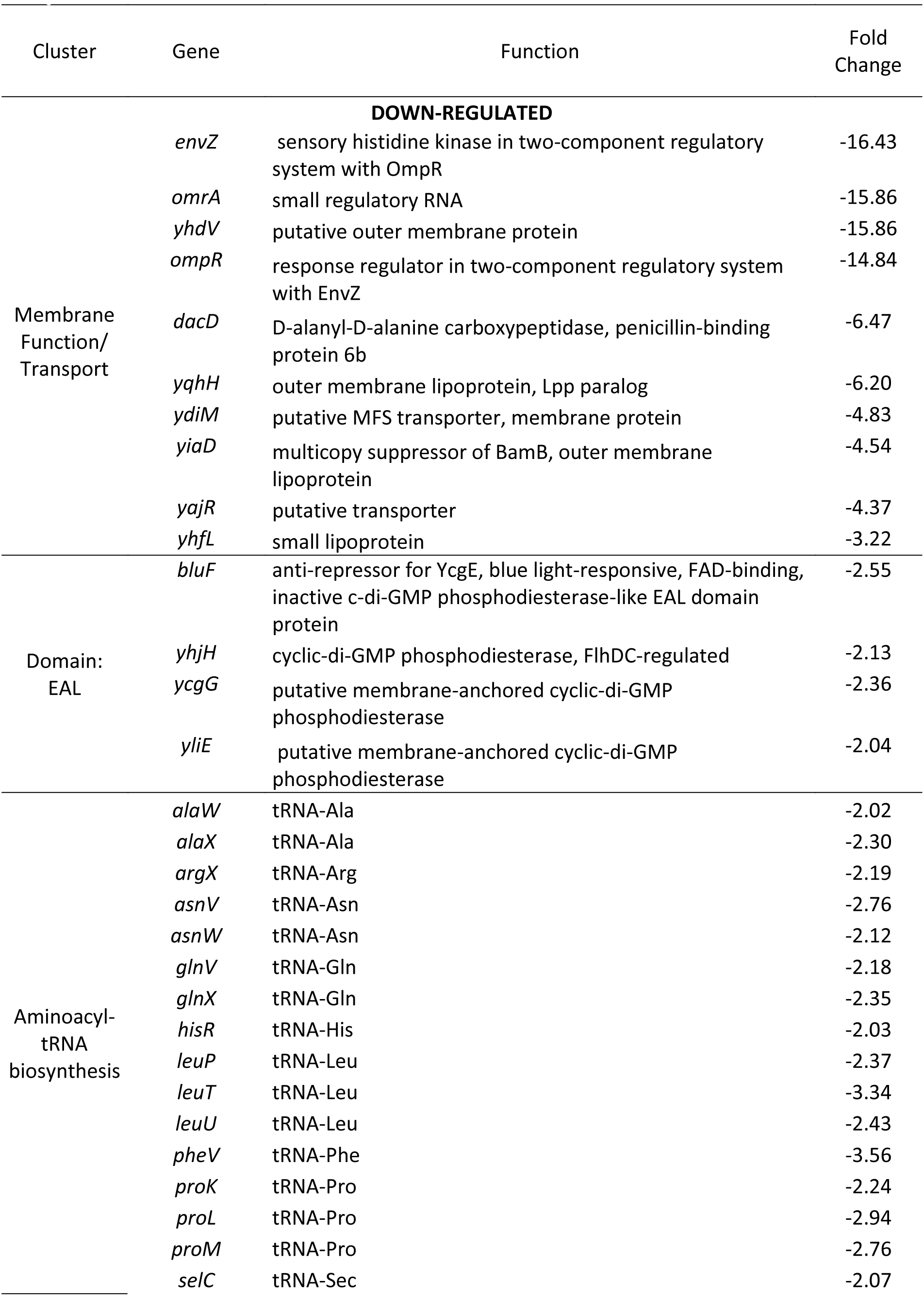
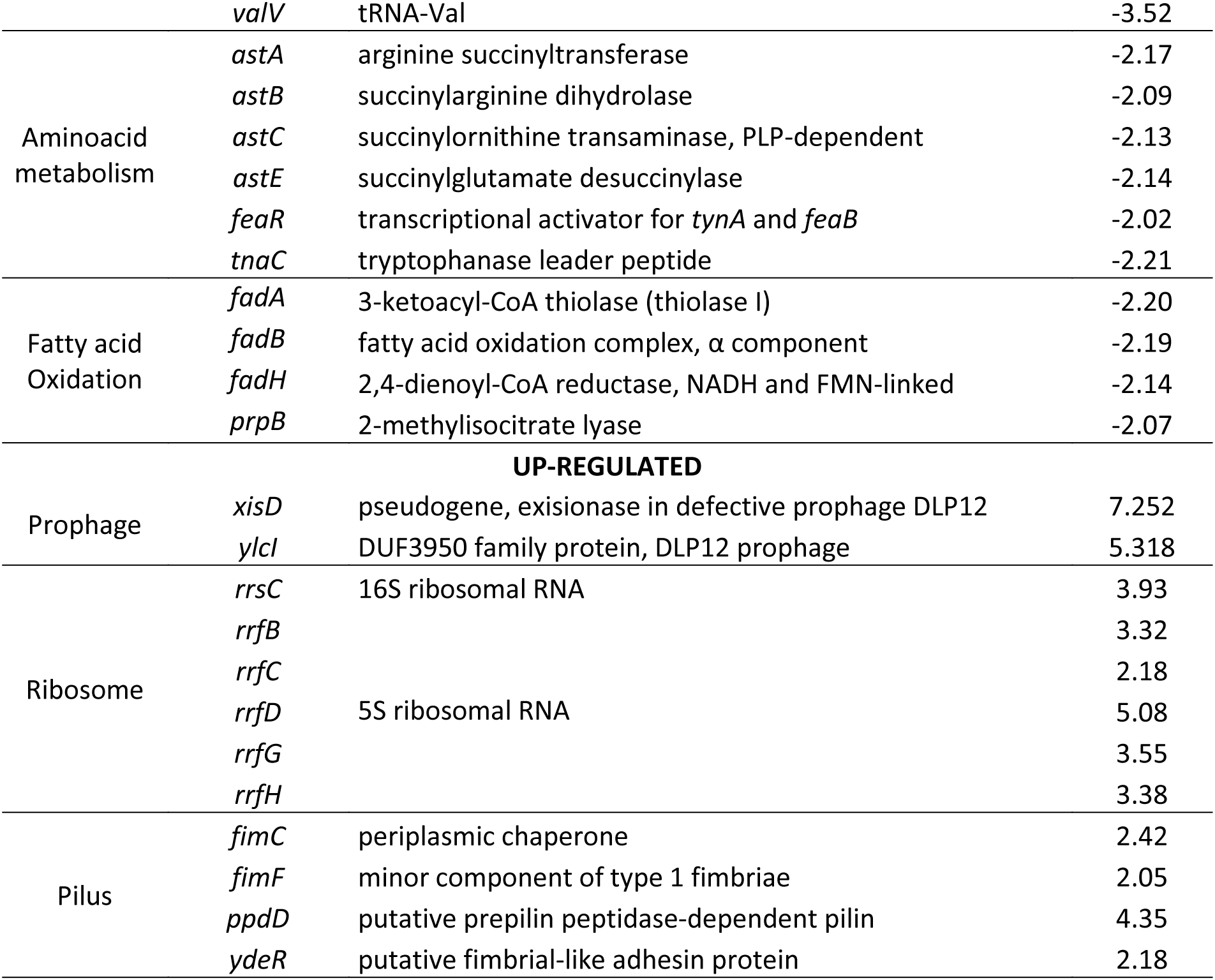
Clustering of the differentially down/up-regulated genes in ARM9 compared to ASD19. Only fold changes >2.0 or <-2.0 are shown.

| Cluster | Gene | Function | Fold Change |
| --- | --- | --- | --- |
| **DOWN-REGULATED** |  |  |  |
| Membrane Function/Transport | *envZ* | sensory histidine kinase in two-component regulatory system with OmpR | -16.43 |
|  | *omrA* | small regulatory RNA | -15.86 |
|  | *yhdV* | putative outer membrane protein | -15.86 |
|  | *ompR* | response regulator in two-component regulatory system with EnvZ | -14.84 |
|  | *dacD* | D-alanyl-D-alanine carboxypeptidase, penicillin-binding protein 6b | -6.47 |
|  | *yqhH* | outer membrane lipoprotein, Lpp paralog | -6.20 |
|  | *ydiM* | putative MFS transporter, membrane protein | -4.83 |
|  | *yiaD* | multicopy suppressor of BamB, outer membrane lipoprotein | -4.54 |
|  | *yajR* | putative transporter | -4.37 |
|  | *yhfL* | small lipoprotein | -3.22 |
| Domain: EAL | *bluF* | anti-repressor for YcgE, blue light-responsive, FAD-binding, inactive c-di-GMP phosphodiesterase-like EAL domain protein | -2.55 |
|  | *yhjH* | cyclic-di-GMP phosphodiesterase, FlhDC-regulated | -2.13 |
|  | *ycgG* | putative membrane-anchored cyclic-di-GMP phosphodiesterase | -2.36 |
|  | *yliE* | putative membrane-anchored cyclic-di-GMP phosphodiesterase | -2.04 |
| Aminoacyl-tRNA biosynthesis | *alaW* | tRNA-Ala | -2.02 |
|  | *alaX* | tRNA-Ala | -2.30 |
|  | *argX* | tRNA-Arg | -2.19 |
|  | *asnV* | tRNA-Asn | -2.76 |
|  | *asnW* | tRNA-Asn | -2.12 |
|  | *glnV* | tRNA-Gln | -2.18 |
|  | *glnX* | tRNA-Gln | -2.35 |
|  | *hisR* | tRNA-His | -2.03 |
|  | *leuP* | tRNA-Leu | -2.37 |
|  | *leuT* | tRNA-Leu | -3.34 |
|  | *leuU* | tRNA-Leu | -2.43 |
|  | *pheV* | tRNA-Phe | -3.56 |
|  | *proK* | tRNA-Pro | -2.24 |
|  | *proL* | tRNA-Pro | -2.94 |
|  | *proM* | tRNA-Pro | -2.76 |
|  | *selC* | tRNA-Sec | -2.07 |
|  | *valV* | tRNA-Val | -3.52 |
| Aminoacid metabolism | *astA* | arginine succinyltransferase | -2.17 |
|  | *astB* | succinylarginine dihydrolase | -2.09 |
|  | *astC* | succinylornithine transaminase, PLP-dependent | -2.13 |
|  | *astE* | succinylglutamate desuccinylase | -2.14 |
|  | *feaR* | transcriptional activator for *tynA* and *feaB* | -2.02 |
|  | *tnaC* | tryptophanase leader peptide | -2.21 |
| Fatty acid Oxidation | *fadA* | 3-ketoacyl-CoA thiolase (thiolase I) | -2.20 |
| | *fadB* | fatty acid oxidation complex, $\alpha$ component | -2.19 |
|  | *fadH* | 2,4-dienoyl-CoA reductase, NADH and FMN-linked | -2.14 |
|  | *prpB* | 2-methylisocitrate lyase | -2.07 |
| **UP-REGULATED** |  |  |  |
| Prophage | *xisD* | pseudogene, exisionase in defective prophage DLP12 | 7.252 |
|  | *ylcI* | DUF3950 family protein, DLP12 prophage | 5.318 |
| Ribosome | *rrsC* | 16S ribosomal RNA | 3.93 |
|  | *rrfB* |  | 3.32 |
|  | *rrfC* |  | 2.18 |
|  | *rrfD* | 5S ribosomal RNA | 5.08 |
|  | *rrfG* |  | 3.55 |
|  | *rrfH* |  | 3.38 |
| Pilus | *fimC* | periplasmic chaperone | 2.42 |
|  | *fimF* | minor component of type 1 fimbriae | 2.05 |
|  | *ppdD* | putative prepilin peptidase-dependent pilin | 4.35 |
|  | *ydeR* | putative fimbrial-like adhesin protein | 2.18 |

To confirm that indeed the reduced transcription of *ompR-envZ* was the major cause for the acquired ammonia resistance, *E. coli* mutant ARM9 was genetically complemented by the introduction of constructs from the ASKA collection(Kitagawa et al 2005) expressing either *ompR* or *envZ*. Introduction of constructs expressing either *ompR* or *envZ* restored ammonia sensitivity, while transformants harboring the empty plasmid continued to be resistant (Figure 5C). This strongly suggests that the reduced expression of *ompR* and *envZ* was the sole cause of the acquired ammonia resistance. It is important to note that mutant ARM9 had also become resistant to AMVCs produced by *S. venezuelae* and by *S. griseus* (the latter grown on glycine), again providing evidence that all strains act by producing ammonia as the AMVC (Figure S3B).

Taken together, these data show that *E. coli* responds to exposure to ammonia by reducing *ompR-envZ* transcription, down regulating the expression of OMPs to minimize the passage of small molecules, and by the reduction of ammonia biosynthesis. Both responses are aimed at defense against the accumulation of toxic levels of ammonia. When exposed to ammonia, *E. coli ompR* mutants were shown to be significantly more sensitive to tetracycline than the parental strain (Bernier et al 2011). Our results show that reducing the expression of OMPs is a defense mechanism against ammonia toxicity extending also earlier observations that *ompF* mutants show impaired response to VOCs that affect the motility of *E. coli* (Kim et al 2013).

### Ammonia released by *Streptomyces* modifies sensitivity to canonical antibiotics

Since ammonia is an AMVC that can reach far from the colony, we considered that the molecule may play a role in long-distance competition with other microbes in the soil, *e.g.* by modifying the effect of other antibiotics produced by actinomycetes, such as erythromycin, kanamycin, actinomycin, spectinomycin and streptomycin. This could be an interesting synergistic effect whereby weapons produced by the strain itself are potentiating via ammonia. To have an indication, the streptomycetes were grown on the left side for 4 days to allow accumulation of compounds on the receiver side containing LB. After that, *B. subtilis* and *E. coli* BREL606 (more resistant to AMVCs) were plated next to *Streptomyces* strains and a filter disk placed on the agar containing different antibiotics. Interestingly, we noticed a significant increase in the sensitivity of *B. subtilis* and *E. coli* to most antibiotics when ammonia-producing streptomycetes were grown adjacent to the receiver cells (Figure 6). Thus, bioactive VCs from *Streptomyces* modulate the activity of soluble antibiotics at longer distances, thereby allowing them to exploit antibiotics produced by other bacteria, or to enhance the activity of its own antibiotics. This novel concept should be worked out further and tested in ecological settings *in situ*, such as competition experiments in controlled microbial communities.

**Figure 6.**
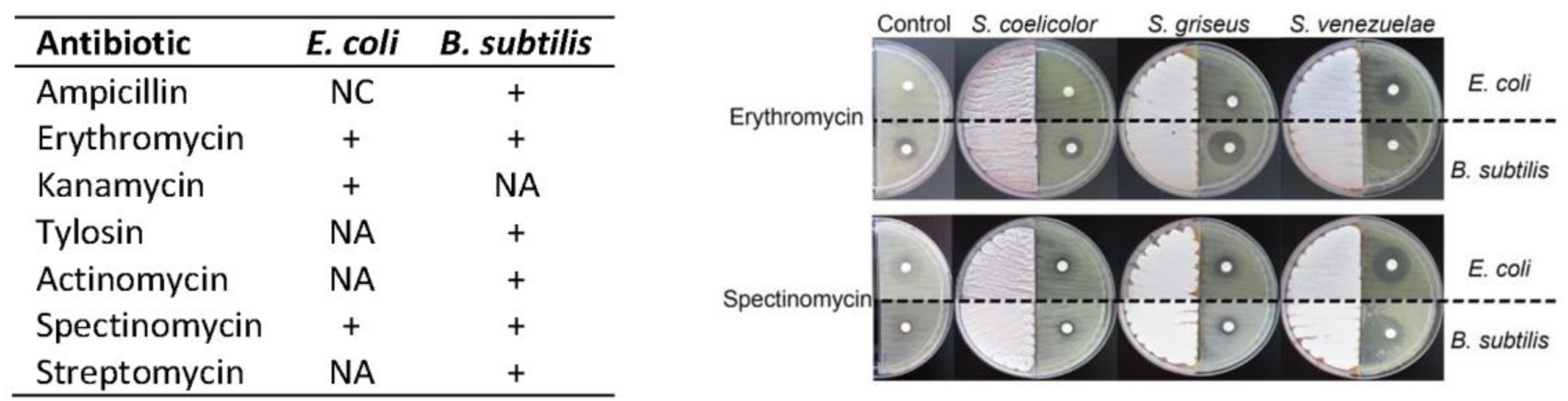
Cooperativity between VCs and soluble antibiotics produced by streptomycetes. Left: changes in antibiotic sensitivity caused by the presence of *Streptomyces* VCs. values indicated increase (+) in halo size. NC, no changes; NA, not active. Right: representative images showing changes in halo size. Streptomycetes were grown on SFM agar, allowed to growth for 4 days, prior to streaking of the indicator strains. Filter disks containing the antibiotic were placed immediately after plating the indicator strains.

### implications on ecology and antibiotic activity

*Streptomyces* VCs have a perceivable impact on the pH of their surroundings. Research has shown that richness and diversity of bacterial soil microbiomes is largely explained by the soil pH(Fierer and Jackson 2006) with acidic soils having the lowest diversity. Basic environments favor bacterial growth while acidic environments do it for fungi(Bárcenas-Moreno et al 2011, Rousk et al 2009). Studies have shown that VCs are more strongly adsorbed in alkaline soils, especially those containing a high organic carbon content (2.9%)(Serrano and Gallego 2006). The release of ammonia by bacteria could have several major implications. First of all, the volatile characteristic allows it to travel far from the producer mediating long-distance interactions. To the best of our knowledge this is the first report showing that soil bacteria such as *Streptomyces* actively kill other bacteria through the air, via the production of ammonia. Additional ecological impact is provided by work of others that shows that ammonia produced by actinobacteria and in particular *Streptomyces* acts as a plant-growth promoter(Passari et al 2017). We hypothesize that such plant-growth promotion may at, least in part, be due to protection against plant pathogenic microbes and in return expand the ‘living room’ of the volatile-producing *Streptomyces*.

Finally, the release of ammonia could help the solubility and diffusion of other types of secondary metabolites, while sensitizing competing bacteria, therefore, the production of a small low-cost ammonia is also a logical strategy to enhance the activity of more complex and hence costly antibiotics, such as polyketides, non-ribosomal peptides or β-lactams. After all, synthesis of these compounds requires expensive high-energy precursors like ATP, NADPH and acyl-CoAs. This is applicable both to antibiotics produced by the organism itself, and to those produced by bacteria further away from the colony. The validity of this concept of “antibiotic piracy” requires further experimental testing.

In conclusion, our work shows that several streptomycetes use ammonia as a low-cost airborne weapon to change their surrounding environment, thereby making their own more costly defense mechanism more effective. In this microbial warfare, the surrounding bacteria then respond by reducing the permeability of their outer membrane and by switching off ammonia production. The field of AMVCs should continue to be studied as it may offer new opportunities for agricultural and/or medical applications, such as for crop protection, plant-growth promotion, and antimicrobial compounds and to continue to understand the role of these molecules in microbial interactions.

## ACKNOWLEDGEMENTS

This work was supported by Grant No. 313599 from The Mexican National Council of Science and Technology (CONACYT) to MA, by VIDI grant 864.11.015 from the Netherlands Organization for Scientific Research (NWO) to PG and by grant 14221 from the Netherlands Organization for Scientific Research (NWO) to GPvW. We thank Hans Zweer for technical help with GC/Q-TOF analysis and Lisanne Storm for the help with the volatile antimicrobial screening, Yasuo Ohnishi and Le Zhang for sharing the glycine cleavage system mutants from *S. griseus* and *S. coelicolor* respectively, and Stephen Douthwaite for providing *E. coli* AS19-RlmA^-^.

